# Comparison of selective genotyping strategies in genomic selection programs for broilers using stochastic simulation

**DOI:** 10.1101/2022.04.29.488103

**Authors:** Setegn. W. Alemu, Anders C. Sørensen, Lei Wang, John Henshall, Rachel Hawken, Per Madsen, Just Jensen

## Abstract

Breeding programs using genomic information have become common in broilers. In broilers, the main traits for selection are body weight and feed efficiency. These traits are measured in both sexes and before sexual maturity. Thus, increases in genetic gain in broiler breeding programs due to the use of genomic information are attributable primarily to the increased accuracy of predicted breeding values of potential parents. As not all birds can be genotyped due to economic constraints, an optimal genotyping strategy needs to be chosen. The objective of this study was to investigate the rates of genetic gain and inbreeding obtained by varying three factors: the percentage of birds genotyped (PG), the percentage of birds genotyped randomly (PRG) or selectively, and the percentage of males among genotyped birds (PMG). Stochastic computer simulation with a full factorial experimental design was used. Significant interaction among the factors (PG, PRG, and PMG) was observed for the rate of genetic gain in body weight and aggregate breeding value, but not for the gain in residual feed intake or the rate of inbreeding per generation. Our findings indicate that the PG, PRG, and PMG need to be considered when selecting a genotyping strategy for a broiler breeding program. If available resources allow only to genotype a small percentage (e.g., 2.5% or 5%) of all birds, the genotyping of 50–100% of male birds and selection of birds to be genotyped using phenotypic information is best. If resources allow to genotype more (e.g., ≥20%) candidates, genotyping of equal numbers of each sex, and low PRG level (i.e., preselection of the majority of candidate based on performance) is best. Provided that a proper genotyping strategy is chosen, we conclude that the incorporation of genomic information in broiler breeding programs can substantially increase the rate of genetic gain.

## Introduction

Genomic selection in animal and plant breeding is the application of dense genetic markers in a breeding program. This approach involves genomic prediction of the genetic merit [genomic estimated breeding value (GEBV)] of individuals based on high-density genetic markers covering the whole genome (Meuwissen et al., 2001). Genomic selection can increase the genetic gain compared with traditional pedigree selection due to the increased accuracy of GEBVs (González-Recio et al., 2009; Daetwyler et al., 2010), and its application can reduce the generation interval, especially when progeny testing is used (Schaeffer, 2006). Furthermore, genomic selection may reduce the rate of inbreeding per generation because it provides additional information about the Mendelian sampling terms of selection candidates (Daetwyler et al., 2007; Dekkers, 2007). Thus, breeding program with genomic information may be a better breeding strategy than breeding program based exclusively on pedigree data.

Breeding programs aim to produce animals with better performance in future generations compared with the current generation. The use of genomic information in these programs has become common for broilers, following similar developments as for dairy cattle and pigs (Wolc et al., 2015). For dairy cattle, where the most important trait is milk yield, genomic information increases the genetic gain by reducing the generation interval. This is because it can be used to predict the breeding value immediately after birth for both sexes, although this trait is recorded after sexual maturity and only for females. In contrast, important traits of broilers, such as growth and feed efficiency, can be recorded before sexual maturity and measured in both sexes. Thus, the accuracy predicting breeding values in broiler breeding programs should be increased by the combined use of phenotypic information and information about pedigree and genomic relationships.

Investigation of the potential contributions of genomic information for broilers using real data may reflect the complexity of the breeding program, but the performance of experiments using different strategies for genomic selection on a commercial scale is costly and time consuming. Furthermore, the number of replicates used in such experiments is often small and non-random. The use of a few replicates or a single replicate renders the testing of several hypotheses about the optimal design of a breeding program difficult (Daetwyler et al., 2013). A good and cost-effective alternative is the use of stochastic simulation. This approach enables investigation of the importance of different design decisions through the simulation of a breeding scheme that resembles real broiler breeding programs as closely as possible. In addition, such models allow for the detailed study of interactions between different factors. However, the use of stochastic models often involves very tedious computation. For example, the examination of six factors at five levels involves the analysis of 5^6^ combinations for the assessment of all possible interactions between factors. Thus, important factors that most influence breeding program outcomes need to be prioritized. Accordingly, we prioritized the percentage of genotyped birds (PG), percentage of randomly chosen genotyped birds (PRG), and percentage of males among genotyped birds (PMG) per time step (*t*) for investigation in this study. Thus, this study was conducted to investigate the importance of PG, PRG, and PMG in each selection round. We compared the rates of genetic gain and inbreeding obtained with the use of different genotyping strategies by conducting stochastic simulation experiments with a full factorial design and the inclusion of PG, PRG, and PMG as main factors. This design enabled the testing of all two- and three-factor interactions between and among the main effects. Arguments for these choices are presented below.

The efficiency of a broiler selection program that incorporates genomic information depends on the number of birds genotyped and phenotyped. The main trait for broilers is the body weight (BW), and this information is typically available for all individuals. However, genomic information is not available for all individuals mainly due to limited availability of genotyping resources. Thus, optimization of the PG is important to maximize the return on genotyping investment. The optimal PG is neither too low (e.g., 1%) nor too high (e.g., 100%). If the PG is too low, the genetic gain generated due to the use of genomic information may not offset the genotyping cost; a comparable rate of genetic gain may be generated without the use of genomic information in this case. The genotyping of a large percentage of candidates may not be necessary because the accuracy of the GEBV increases and the genetic gain diminishes with increasing PG (Goddard and Hayes, 2009). In other words, at some PG, the marginal return derived from the genotyping of an additional animal is less than the marginal genotyping cost (Henryon et al., 2014). Thus, investigation of how the PG affects the genetic response in a breeding program is essential.

The second factor we considered in this study was the PGR. Data on economically important broiler traits, such as the BW and feed efficiency, are usually available in the form of performance and/or pedigree information. They can be used as selection criteria for genotyping because birds with the best performance are expected, on average, to have the largest GEBVs and thus a high probability of being chosen as parents (Henryon et al., 2014). However, when the PG is large, the genotyping of a smaller proportion (<25%) of candidates randomly and the remaining candidates based on phenotypic information may generate a similar rate of genetic gain as would the genotyping of all candidates based on phenotypic performance information. The genotyping of a random selection of candidates also helps to reduce the bias in the estimation of variance components when these components are unknown and genomic selection is practiced for several generation**s** (Wang et al., 2020). Thus, investigation of the rates of genetic gain obtained by different PRGs is essential.

The next natural question following on consideration of the PG and PRG regards the distribution of female and male genotyped individuals. In dairy cattle, the largest benefit originates from the genotyping of sires, which are progeny and tested with high accuracy. The inclusion of females in the genotyped population is essential only when the population is small (Mc Hugh et al., 2011; Thomasen et al., 2014, 2016; Sollero et al., 2019). The optimal percentages of male and female genotyped animals may not be similar for broilers to those for dairy cattle because broiler sires have fewer offspring and dams have more offspring than do dairy cattle sires and dams. Furthermore, traits are recorded in both sexes and before sexual maturity for broilers. Thus, a training population including both sexes of broiler may be better than a single-sex population. For example, using real data, Lourenco et al. (2015) demonstrated that the use of a two-sex broiler training population increased the selection accuracy relative to the use of single-sex populations. Apart from that study, no study to our knowledge has been conducted to determine the optimal percentage of males in a broiler training population or how this percentage interacts with other parameters determining the population. Thus, investigation of the rates of genetic gain obtained by different PMGs is essential.

## Methods

Stochastic simulation was used to estimate direct and correlated responses to selection and the rate of inbreeding. The simulation was done in three phases termed historical phase, pedigree–best linear unbiased prediction (BLUP), and genomic phase. In the historical phase, a population with non-overlapping generations was simulated forward in time using QMSIM software (Sargolzaei and Schenkel, 2009). In total, 950 non-overlapping generations with a gradual linear increase in population size from 1000 to 2400 individuals per generation were simulated to generate initial linkage disequilibrium (LD) and mutation-drift balance. The numbers of male and female individuals were equal throughout this phase, except in the last historical generation, which consisted of 20 males and 2380 females. We chose this historical phase to generate an LD structure similar to that observed in empirical broiler data (Figure 1). The mating system in this phase consisted of the union of gametes drawn randomly from the male and female pools.

**Figure 1.**
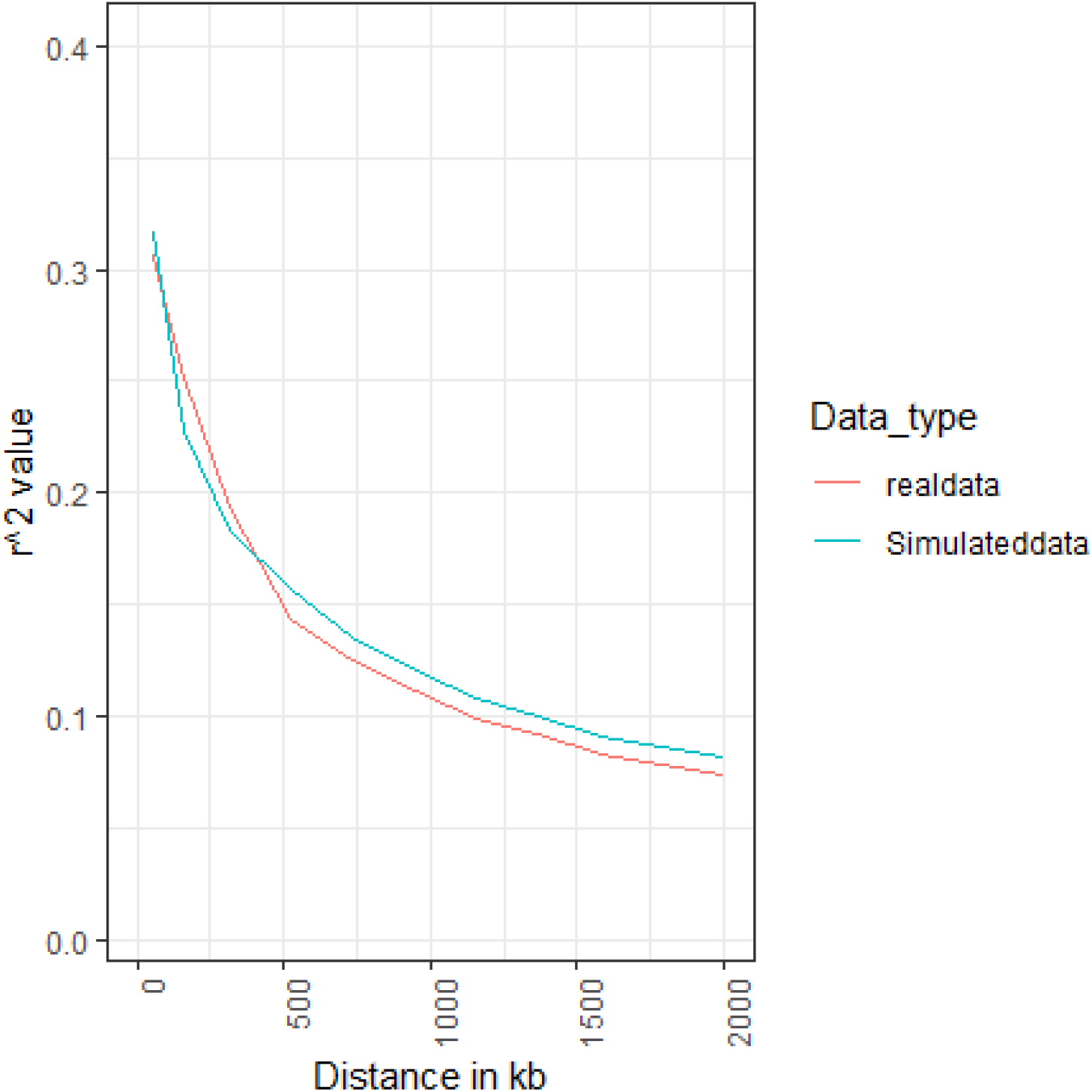
Linkage disequilibrium decay in terms of correlation between markers of increasing distance for real (Wang et al., 2020) and simulated data.

The pedigree-BLUP and genomic phases lasted for 20 timesteps (*t*s) each (1–20 and 21–40. respectively). In the genomic phase, a subset of individuals was genotyped in each *t*, depending on the scenario investigated (PM, PRG, or PMG). In each *t*, birds were hatched and a selection process (trait recording, breeding value estimation, and selection of parents) took place. Generations were overlapping, and each generation corresponded to 6.5 *t*s. The second and third phases of the breeding programs studied were simulated using the ADAM software (Pedersen et al., 2009).

### Genome

The simulated genome consisted of 26 chromosomes with a total length of 916 cM and individual chromosome lengths resembling those in the broiler genome (Burt, 2005). In the first historical generation, the number of marker and quantitative trait locus (QTL) alleles at each locus was 2, with equal frequency (0.5). A recurrent mutation rate of 2 *x* 10^-5^ was considered for markers and QTLs to establish mutation-drift equilibrium in the historical generations. Thus, mutation and drift were the only evolutionary forces that existed in this phase. To achieve the desired number of segregating loci after 950 generations, about 17 times as many biallelic loci as needed were simulated. In the last historical generation, 40K single nucleotide polymorphisms (SNPs) and 2K QTLs were sampled randomly to build an SNP panel and define the QTL effects. For quality control, SNPs and QTLs with minor allele frequencies ≤ 5% were excluded.

### Traits simulated

The breeding goal consisted of two traits: the BW and residual feed intake (RFI). In the pedigree-BLUP and genomic phases, the BW phenotype was measured for all individuals and the RFI phenotype was measured for the 20% of heaviest males and 20% of heaviest females in each *t*. Both phenotypes were available when the birds were one *t* old. The genetic parameters used for the simulation are given in Table 1 and were chosen from previous studies of broiler BW and RFI (Aggrey et al., 2010; Begli et al., 2016).

**Table 1.**
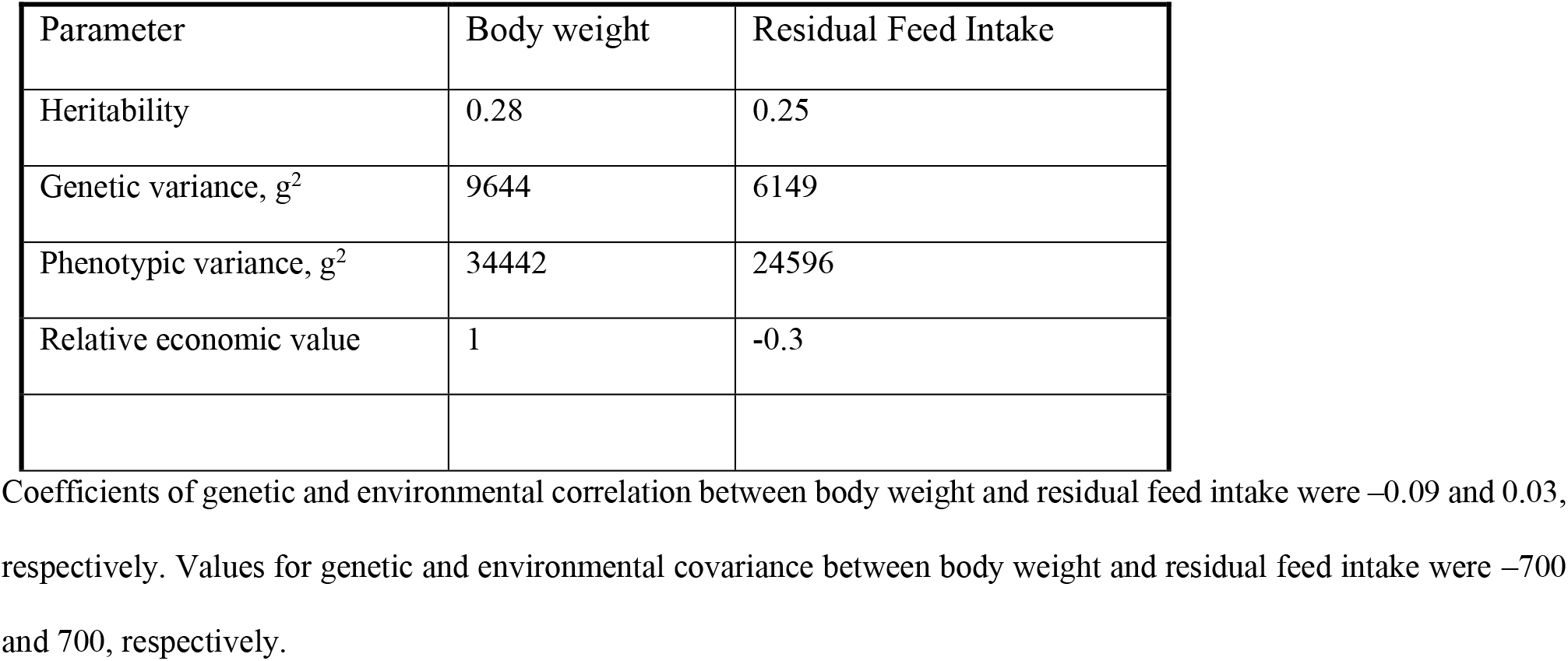
Genetic parameters used for the simulation.

| Parameter | Body weight | Residual Feed Intake |
| --- | --- | --- |
| Heritability | 0.28 | 0.25 |
| Genetic variance, $g^2$ | 9644 | 6149 |
| Phenotypic variance, $g^2$ | 34442 | 24596 |
| Relative economic value | 1 | -0.3 |

Coefficients of genetic and environmental correlation between body weight and residual feed intake were $-0.09$ and $0.03$ , respectively. Values for genetic and environmental covariance between body weight and residual feed intake were $-700$ and $700$ , respectively.

### Aggregate breeding value

The aggregate genotype is a linear index: *H* = *v_1_g_1_* + *v_2_g_2_*, where *H* is the aggregate breeding value, *v_1_* and *v_2_* are the partial economic weights given to the BW and RFI, and *g*_1_ and *g_2_* are the breeding values for the BW and RFI, respectively. The economic value of a trait was defined as the effect of a marginal (one-unit) change in the genetic level of the trait on the objective function (i.e., profit), with all other traits included in the aggregate genotype kept constant. We used an economic weight for the BW that was 3.3 times that for the RFI. As the BW and RFI are phenotypically independent, the value of 1 g live chicken is the same as the value of 3.3 g feed. Thus, we chose economic values of 1 for the BW and −0.3 for the RFI. The negative RFI value indicates that the objective is to reduce the RFI.

### Breeding value

The additive allelic effect of each QTL was drawn independently from a normal bivariate distribution. True breeding values were simulated by adding the QTL allelic effects within and across loci. The BW and RFI phenotypes were simulated by adding the residual term sampled from a bivariate normal distribution, with a mean value zero and residual variance-covariance specified in Table 1, to the true breeding values.

### Population structure

The breeding population for each *t* consisted of 52 sires and 520 dams. Each sire was mated randomly to 10 dams and each dam produced 10 offspring with an equal sex ratio; thus, 5200 birds were hatched in each *t*. Fifty-two half-sib families with a size of 100 and 520 full-sib families with a size of 10 were present in each *t*. Thirteen new sires and 130 new dams were selected in each *t*, when the birds were one *t* old. Individuals selected as parents were parents for four *t*s.

#### Statistical estimation of breeding values

In the pedigree-BLUP phase, additive breeding values were estimated using a bivariate BLUP animal model with a numerator relationship matrix (**A**) serving as the covariance structure (Henderson, 1975). In the genomic phase, breeding values were estimated using a bivariate single-step genomic best linear unbiased prediction (ss-GBLUP) animal model with a relationship matrix that included genomic and pedigree information (**H**) (Legarra et al., 2009; Christensen and Lund, 2010). The formula used to estimate breeding values during these two phases was:

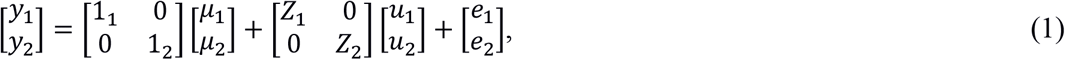

where 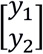 is a vector of BW and RFI phenotypes, 11 and 12 are unit vectors for the population means, 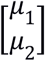 is the vector of population means (intercept) for the BW and RFI, and 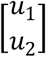 is the vector of breeding values for the BW and RFI. *Z*_1_ and *Z*_2_ are design matrices that associate breeding values with the BW and RFI, and 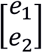 is the vector of the random residuals of BW and RFI.

In the pedigree-BLUP phase, the breeding values were assumed to have a multivariate normal distribution as 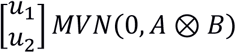, where **A** is the genetic relationship matrix derived from pedigree information and 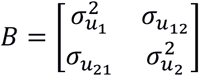 is the variance-covariance matrix of additive breeding values for the BW and RFI. In the genomic phase, **A** was replaced by **H**:

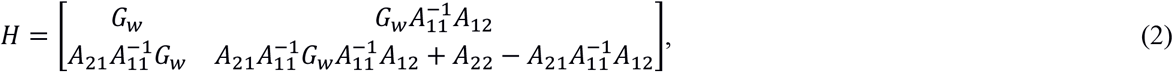

where **A_11_** is the sub-matrix of **A** for genotyped animals, **A_22_** is the sub-matrix of **A** for non-genotyped animals, **A_12_** (or **A_21_**) is the sub-matrix of **A** for relationships between genotyped and non-genotyped animals, and **Gω** = (1 – ω) **G** + ω**A**_11_, where ω is a weight with a value 0.25 and **G** is the genomic relationship matrix (VanRaden, 2008).

The variance of the residuals 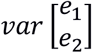 is a block diagonal matrix with two types of diagonal block: one for individuals in which both traits are measured and the variance-covariance matrix is 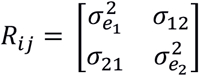, and one for individuals in which only the BW is measured. The (co)variance is 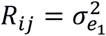. All residual covariances between traits measured in different birds were assumed to be zero. In all rounds of selection, we obtained EBVs or GEBVs by considering all information from *t* = 1 until the current *t*. Breeding values were estimated using the DMU software package (Madsen and Jensen, 2013). Parents were selected based on EBVs in the pedigree-BLUP phase and on GEBVs in the genomic phase. In both phases the goal was to maximize genetic gain in the aggregate breeding value (weighted linear combination of body weight and residual feed intake where the weight is the economic value).

In summary, the simulation we implemented mimics a breeding program similar to a real broiler breeding program. For example, we used genetic parameters from previous studies for both traits, and the LD and population structures were close to those of a real breeding program. Furthermore, we used pedigree-BLUP (the optimal use of pedigree information) to estimate breeding values in the first 20 *t*s and ss-GBLUP (the optimal use of information from genotyped and non-genotyped individuals) in the last 20 *t*s (Legarra et al., 2009; Christensen and Lund, 2010).

### Factors investigated

We investigated the rate of genetic gain obtained with aggregate breeding values, the BW, and the RFI using a simulation with variance of the three factors of interest (PG, PRG, and PMG; Table 2). All combinations of these factors (4 x 5 x 5 = 100) were investigated in a full *in-silico* factorial experiment.

**Table 2.**
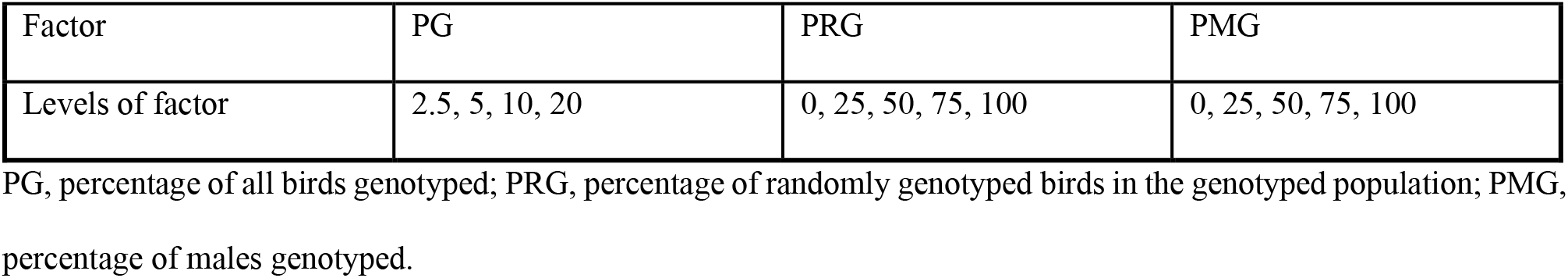
Factors investigated in the study.

#### Analysis of responses to selection and accumulated inbreeding

All 100 scenarios were replicated 50 times. For each replicate, a set of variables (the true breeding value for the aggregate breeding value, BW, RFI, and inbreeding for all birds simulated) was obtained and comprised the simulation output. We summarized this information using the rate of change in the aggregate breeding value (dH) by estimating the coefficient for the regression of the aggregate breeding value on *t*, and performing similar calculations for the BW (dBW), RFI (dRFI), and genomic inbreeding (dF). First, the average of the true breeding value (from 5200 birds in each *t*) for H, the BW, and the RFI from each *t* and the average level of genomic inbreeding (from 5200 birds in each *t*) were computed. Second, we calculated regression coefficients for *t*s 31–40, when the genomic information was well incorporated in the breeding program due to the use of genomic information in the selection rounds 21-30. Most of the birds’ performances for *t*s 31–40 contained the genomic information. To facilitate comparison with findings from other studies, we used the generation instead of the *t* for examination of the rate of genomic inbreeding. Genomic inbreeding values were calculated by subtracting 1 from the diagonal of genomic relationship. The change in the rate of genomic inbreeding per generation was calculated as *1 –exp(β)*, where *β* is a linear regression of l*n* (1 – F*g*) on *g*, and F*g* is the average degree of genomic inbreeding at each *t*.

We performed a three-factor analysis of variance (ANOVA; R Core Team, 2017) of the regression coefficients, which included all main effects (PG, PRG, and PMG), the two factors, and the three-factor interaction. We did not include replicate effects in the analysis because all replicates were simulated independently, with new streams of random numbers used in each simulation. Thus, no relationship existed between any of the 50 replicates.

## Results

The three-factor ANOVA showed that all main effects were highly significant. Table 3 shows the levels of significance for all two- and three-factor interactions between and among PG, PRG, and PMG. The three-factor interaction was significant for dH and dBW, but not for dRFI or dF. All two-factor interactions and each of the factors was significant. In presenting the results, we focus on the three-factor interactions for dH and dBW.

Significance of two- and three-factor interactions.

Significance of two- and three-factor interactions.
| <sup>1</sup>response | PMG*PRG | PMG*PG | PRG*PG | PMG*PRG*PG |
| --- | --- | --- | --- | --- |
| dH | < 0.0001 | < 0.0001 | < 0.0001 | < 0.0001 |
| dBW | < 0.0001 | < 0.0001 | < 0.0001 | < 0.0001 |
| dRFI | < 0.0001 | < 0.0001 | 0.003 | 0.1 |
| dF | < 0.0001 | 0.03 | 0.016 | 0.052 |

<sup>1</sup>All main effects were significant at $p < 0.0001$ .
PRG, percentage of randomly genotyped birds in the genotyped population; PMG, percentage of males genotyped; PG, percentage of all birds genotyped; dH, rate of genetic gain in the aggregate breeding value; dBW, rate of genetic gain in body weight; dRFI, rate of genetic gain in residual feed intake; dF, change in the rate of inbreeding.

### dH

As effects involving three interacting factors can be difficult to grasp, we present the results in interaction plots. Figure 2 shows five interaction plots for dH. The dH increased with the PG in all cases, although the rate of increase depended on the PRG and PMG. The dH units per *t* ranged from 18 to 23.5, representing most of the potential advantage derived from the application of genomic information in the breeding program. The standard errors of selection responses ranged from 0.1 to 0.2.

**Figure 2.**
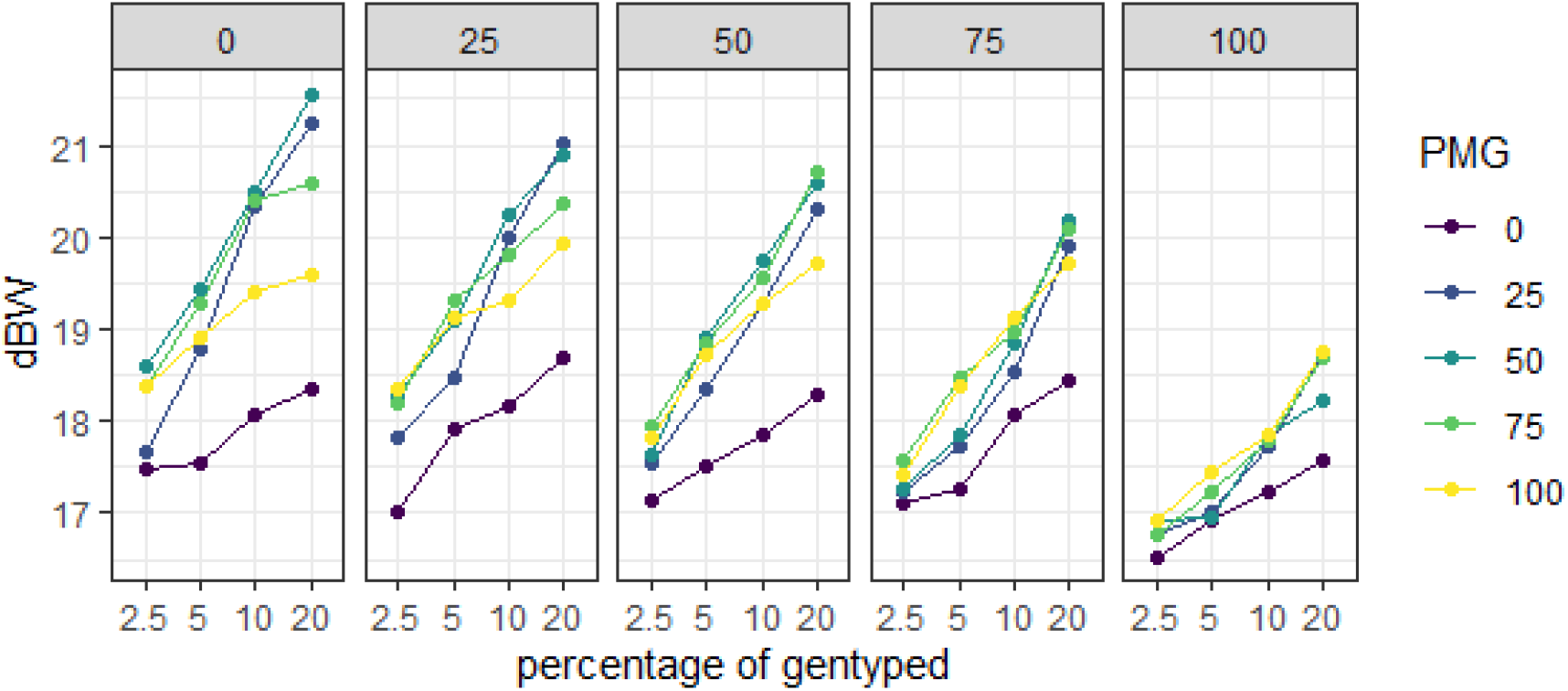
Interaction plots for the genetic gain in the aggregate breeding value (dH) per time step (*t*) for all combinations of the percentage of males genotyped (PMG i.e. color of the plot), percentage of birds genotyped (PG) (X-axis label), and percentage of birds randomly genotyped (PRG)(shown on top of each sub plot).

The positive effect of PG on dH was largest when PRG = 0, i.e., when only selective genotyping was applied, diminished gradually as the PRG increased, and was smallest at PRG = 100 (i.e., all birds to be genotyped were selected at random). The effect of PG on dH also differed according to the PMG; it was smallest at PMG = 0, and larger at PMG = 50 than at PMG = 100. Thus, if birds of only one sex are genotyped, they should be males, but the genotyping of similar numbers of males and females generates a greater genetic gain.

The positive effect of PG on dH depended on the PRG and PMG levels. When both the PRG and PMG were 0, the effect of PG on dH was smaller. With PRG = 0 and the PMG increasing from 0 to 0.25 and 0.5, the positive effect of PG on dH increased; with further increase in the PMG from 0.5 to 0.75 and 1, the positive effect of PG on dH decreased. Thus, PMG and PG interact at PRG = 0 because the positive effect of PG on dH is not constant for different PMG levels. As the PRG increased, the interaction between PMG and PG decreased. In summary, for large and small PG values, selective genotyping should be used, and approximately equal numbers of males and females should be genotyped. The advantage of this approach is substantially greater for larger PG than for smaller PGs.

### dBW

Figure 3 shows five interaction plots for the dBW. For given scenarios, dBW values were slightly smaller than dH values, although the two parameters exhibited similar trends. For example, the dBW at PRG = 0, PMG = 50, and PG = 20 was 21.6, whereas the dH in this scenario was 23.3.

**Figure 3.**
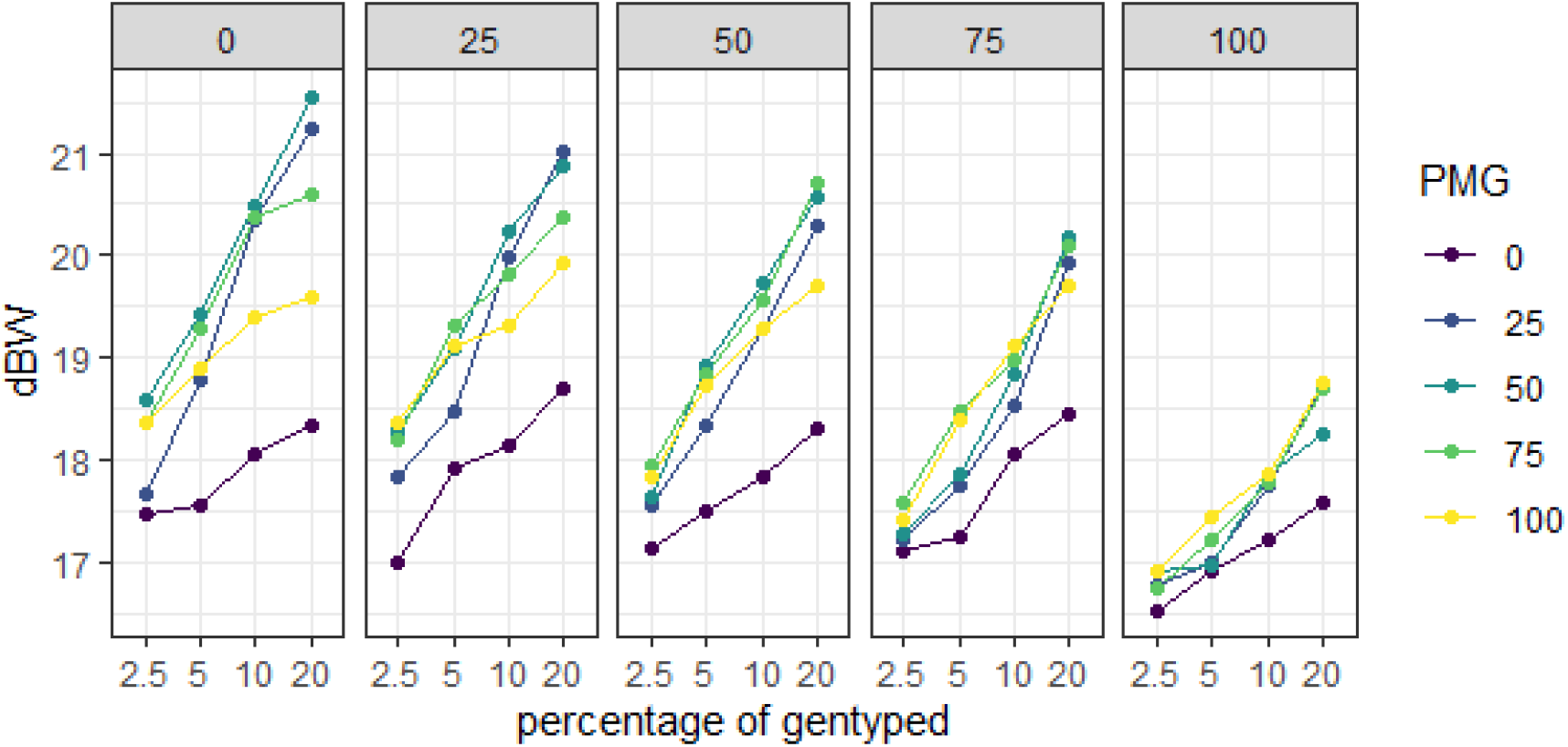
Interaction plots for the genetic gain in body weight (dBW) per time step (*t*) for all combinations of the percentage of males genotyped (PMG i.e. color of the plot), percentage of birds genotyped (PG) (X-axis label), and percentage of birds randomly genotyped (PRG)(shown on top of each sub plot).

### dRFI

Figure 4 shows five interaction plots for the dRFI. dRFI values were small in all scenarios [range, −3.3 to −5.8 g (standard error, 0.15 to 0.19) per *t*] and differed little among scenarios. Unlike the dBW and dH values, the dRFI values exhibited no clear trend as a function of the PG, PMG, or PRG. Furthermore, the rate of genetic gain for dRFI was considerably lesser than the rate of genetic gain for dBW.

**Figure 4.**
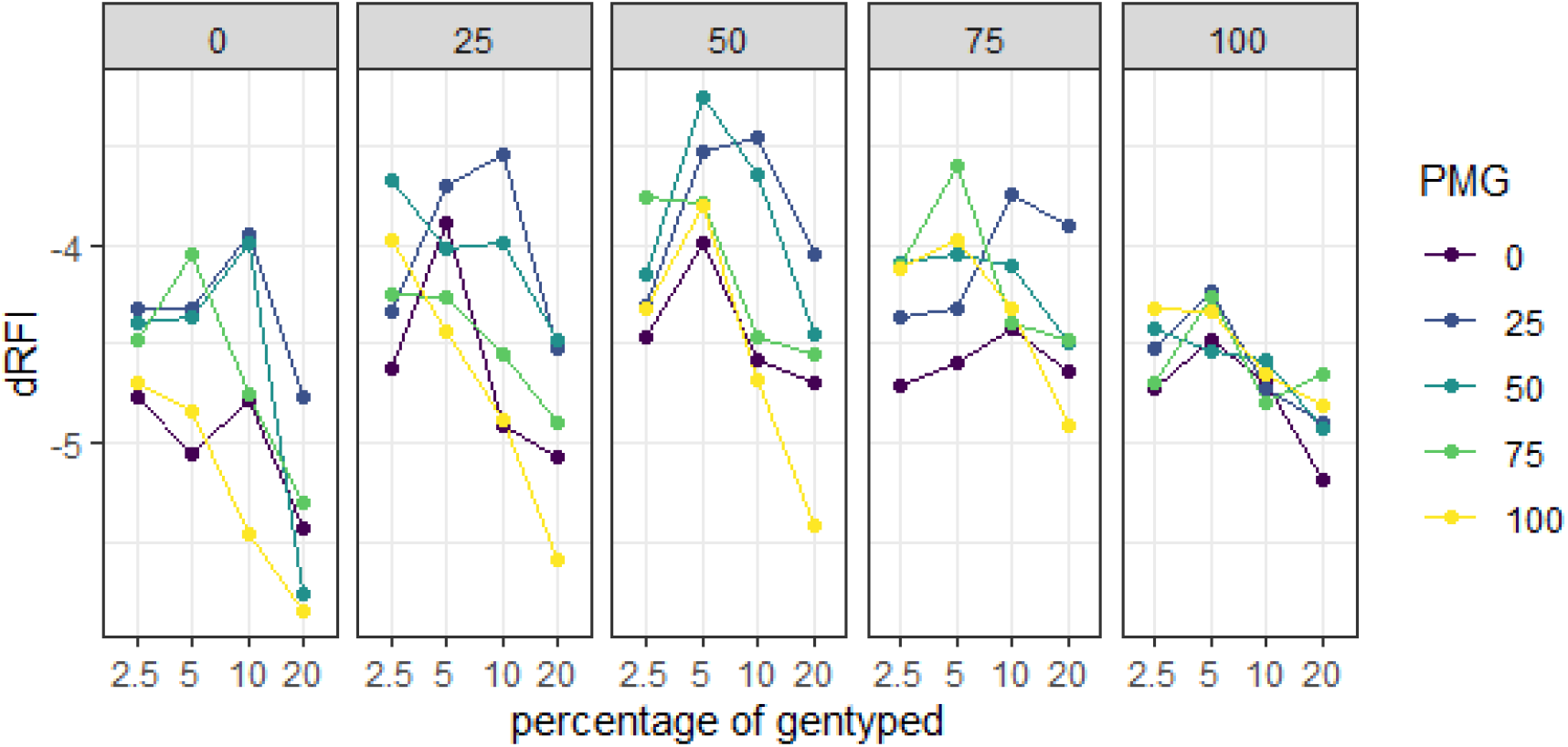
Interaction plots for the genetic gain in residual feed intake (dRFI) per time step (*t*) for all combinations of the percentage of males genotyped (PMG)(color of the plot), percentage of birds genotyped (PG) (X-axis label), and percentage of birds randomly genotyped (PRG)(shown in each sub plot).

### dF

Figure 5 shows five interaction plots for the dF. dF values ranged from 0.01 to 0.02 (standard error, 0.0005–0.001). Although not as clear as for the dH, the dF exhibited two trends: it was larger when only females were genotyped than when only males were genotyped, and larger at PG = 2.5 than at PG = 20.

**Figure 5.**
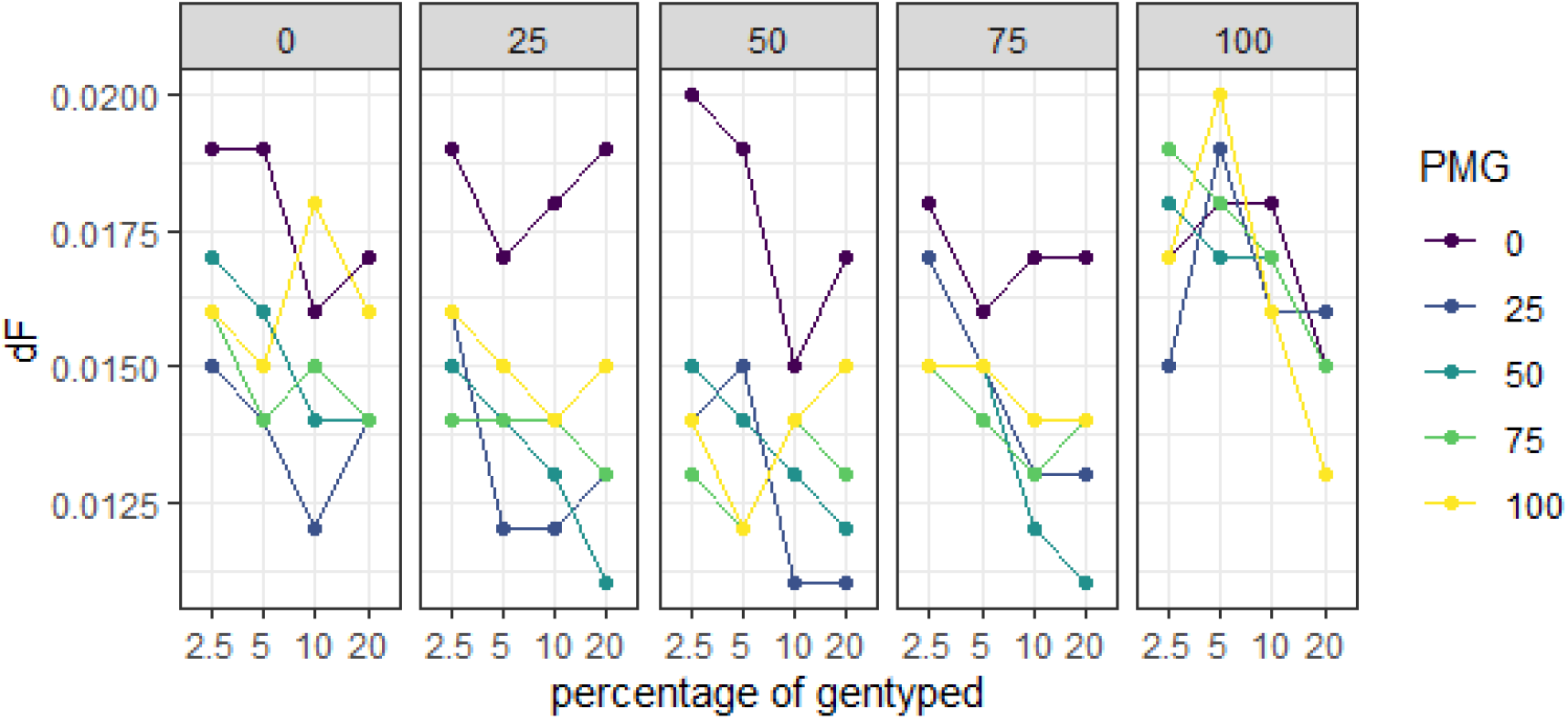
Interaction plots for the rate of inbreeding (dF) per generation for all combinations of the percentage of males genotyped (PMG), percentage of birds genotyped (PG), and percentage of birds randomly genotyped (PRG).

Comprehensive results for all 100 scenarios are available as three different views created with three graphical tools (ggplot, ploty, and Lattice plot) are available via the shiny app link https://setwork.shinyapps.io/Broiler_genomic_selection/. The source code used to produce the results is available via the GitHub link https://github.com/setegnworku/Genomic-selection-in-broiler.

## Discussion

In this study, we investigated the effects of the PG, PRG with selection of the remaining genotyped birds according to the largest BWs among contemporaries, and the PMG per *t* in broiler breeding programs. All factors were expected to influence the rates of genetic gain and inbreeding in programs incorporating genomic information. The dH differed considerably with different PMG and PG levels across PRG levels. Thus, it was affected substantially by interactions between PG, PRG, and PMG, and these three factors need to be considered simultaneously when choosing a genotyping strategy for a broiler breeding program.

### dH

dH values generally increased with the PG due to the greater accuracy obtained by genotyping (i.e., the accuracy of selection based on breeding values that incorporate genomic information increases with the PG) (Daetwyler et al., 2008; Goddard 2009). Larger PGs were beneficial, although a clear diminishing effect was observed from PG = 2.5 to PG = 20. We thus investigated the effects of PG = 40 and PG = 50 to validate our hypothesis, diminishing effect. In consideration of the computation time, we performed this investigation for a single scenario with equal proportions of sexes and the heaviest selection candidates genotyped. The results are shown in Figure 6.

**Figure 6.**
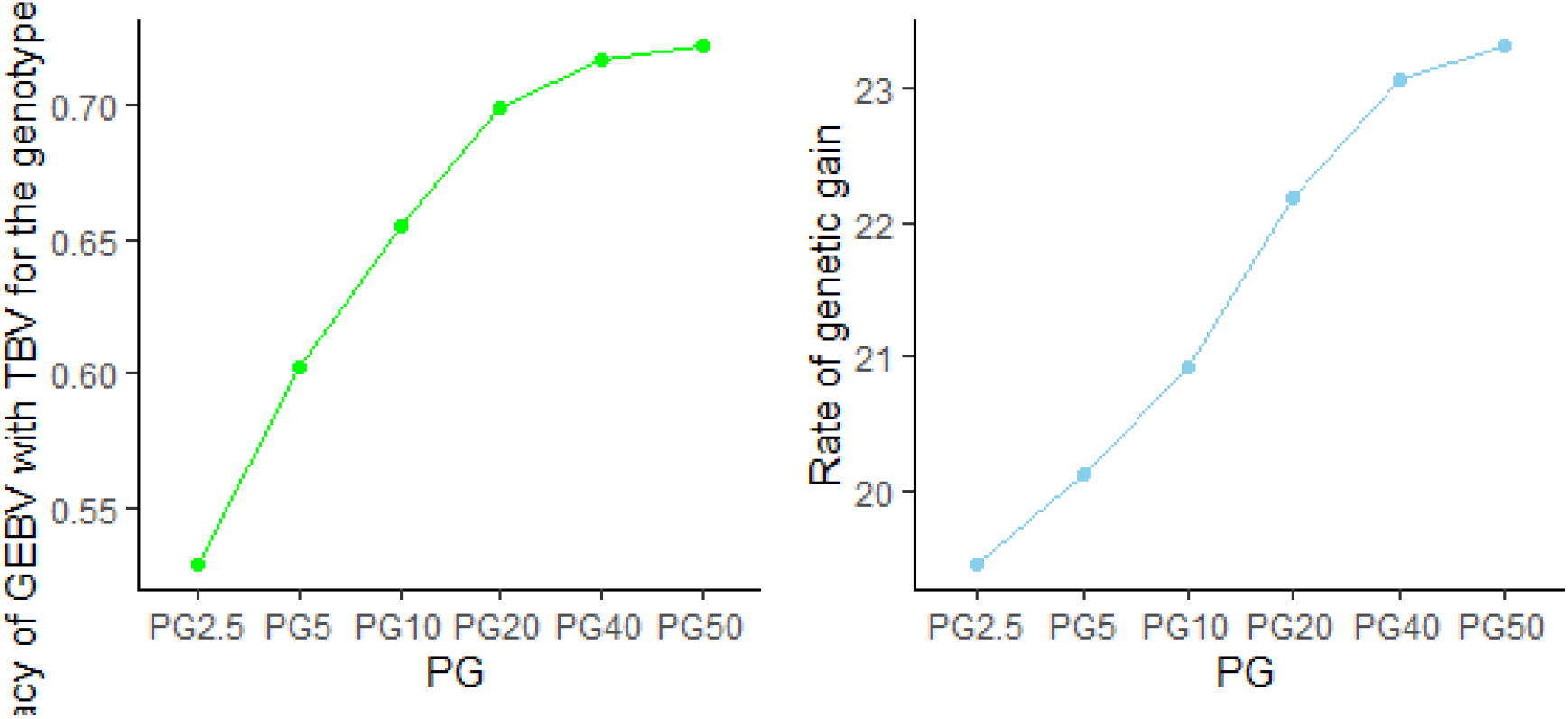
Accuracy of selection [correlation between the genomic estimated breeding value (GEBV) and true breeding value (TBV) for selection candidates] and rate of genetic gain when equal proportions of sexes (PMG = 50) and the heaviest selection candidates (PRG = 0) are genotyped.

The dH increase diminished toward 0 as the PG approached 50. Whereas an increase in the PG from 10 to 20 increased the genetic gain by 6% and an increase from 20 to 40 increased the genetic gain by 4.4% (2.2% per 10% change), an increase in the PG from 40 to 50 increased the genetic gain by only 1.2% (i.e., the rate of genetic gain was approaching a plateau). A further increase in the PG (e.g., to 60), would increase the genetic gain relative to PG = 50 by less than 1%. The accuracy of selection showed a similar trend; for example, it increased by 1.4% (from 0.71 to 0.72) when the PG changed from 40 to 50. Henryon et al. (2012) reported similar results, showing that the maximum genetic gain can be realized by genotyping 40–50% of the selection candidates when animals to be genotyped are chosen based on phenotypic information.

The positive effect of PG increases was larger at PRG = 0 in this study because the genotyping of many, mainly heavy, birds increase the probability of selecting individuals with high breeding values and accuracy. Phenotypic records of BW have a large influence on predicted breeding values and genotyping of heavy birds helps to distinguish their additive genetic merit. As the PRG increases, the positive effect of the PG decreases because genotyped and non-genotyped birds may be selected as parents. The selection of non-genotyped birds reduces the accuracy of predicted breeding values. Thus, at PRG = 0 the positive effect of the PG is larger because the chance of selecting genotyped individuals is increased, whereas at PRG = 100 the positive effect of the PG is smaller because the chance of selecting genotyped individuals is decreased.

The positive effect of the PG on the dH was larger when only males were genotyped (PMG = 100) than when only females were genotyped (PMG = 0) in this study because the selection intensity is larger for males than for females. Genotyping of more males increases the probability that genotyped individuals are selected as parents, thereby increasing the accuracy of predicted breeding values. This result is in line with Henryon et al.’s (2012) finding that the genetic gain is maximized by genotyping only males at a low PG in pig breeding programs (which have a structure similar to that of broiler programs).

The positive effect of the PG depended on the PRG and PMG levels in this study. It was maximal at PRG = 0 and PMG = 50 because genotyping of both sexes increases the probability that genotyped birds are selected based on breeding values predicted with high accuracy. Thus, the selective genotyping of the heaviest birds increases the genetic gain more with a larger PG than with a smaller PG. This result is in accord with the finding of Henryon et al. (2012) that the genotyping of both sexes at PGs ≥ 20 increased the genetic gain in a stochastic simulation of pig breeding programs. It is also consistent with Lourenco et al.’s (2015) finding in an analysis of real data that the genotyping of both sexes increased the accuracy of selection compared with the genotyping of only one sex.

In this study, a constant percentage of individuals was genotyped during the entire genomic phase in a given scenario. For example, with PG = 20, (1080 x 20=20800) genotyped individuals were redistributed equally (*n* = 1080) in each *t*. Further investigation of whether genotyped individuals can be distributed in another way in different *t*s [e.g., via the genotyping of more (e.g., 1500) birds in *t*s 21–25 and fewer (e.g., 886) birds in *t*s 26–40] to generate more genetic gain than obtained with a constant PG is warranted.

### dBW

The dBW increases with the PG because the training population also increases, given the availability of phenotypic information on BW for all birds. With this increase in the training population, the accuracy of selection based on breeding values that incorporate genomic information also increases (Daetwyler et al., 2008; Goddard, 2009).

The rate of genetic gain in H is dominated by the rate of genetic gain in BW because the BW is economically much more important than the RFI. The two traits have similar heritability, but the genetic variance in the BW is 57% greater than that in the RFI. The RFI was assessed in only 20% of the heaviest birds in this study, and selections for the two traits were nearly independent of each other, as the genetic correlation between them was weak (−0.07).

### dRFI

dRFI values were substantially lower than dBW values in this study, as the economic value of the BW was 3.3 times the per-gram RFI. At least three factors explain this difference. First, the number of individuals with phenotypic RFI information was limited (520 males and 520 females in each *t*), meaning that a large proportion of birds, especially females (*n* = 130), should be chosen from this subset to select for the RFI. In contrast, phenotypic BW information was available for all individuals, increasing the possibility of selection for this trait. In addition, phenotypic and genotypic RFI information was available for fewer individuals than was phenotypic information alone in most scenarios. Second, the BW is slightly more heritable than RFI (0.28 vs. 0.25), and the genetic variance in the BW is 57% greater than that in the RFI. Third, a large number of individuals was phenotyped but not genotyped for the BW compared with the RFI. The use of ss-GBLUP aids the efficient utilization of this information, which increases the rate of genetic gain for the BW.

### dF

dF values were larger at PMG = 0 than at PMG = 100 in this study. When only females are genotyped, males are basically selected on the basis of their own performance combined with pedigree information. Compared with the use of genomic information, the use of pedigree information increases the probability that close relatives will be co-selected because the former provides more information about Mendelian sampling (Daetwyler et al., 2007). At PMG = 100, the probability of co-selection of close female relatives increases because no genomic information is available for females. Thus, the co-selection of male relatives increases dF more than does the co-selection of female relatives because males have more offspring than do females (Robertson, 1961).

dF values were largest with small PG values (2.5 and 5) and decreased at PGs of 10 and 20 due to the increased probability of selecting parents from more families (Daetwyler et al., 2007). At low PGs, the single-step evaluation model is very similar to a pedigree animal model with no birds genotyped. The use of the animal model increases the probability that close relatives will be co-selected. Models with larger PG values contain more information on Mendelian sampling effects, reducing this probability and thus the rate of inbreeding (Daetwyler et al., 2007).

#### Comparison of selected scenario outcomes with pedigree-BLUP results

To investigate the effect of the inclusion of genomic information in a broiler breeding program, we compared the rate of genetic gain per *t* using pedigree-BLUP with selected scenarios (PRG = 0 or 100, PMG = 50, and all PG levels; Table 4). The genetic gain obtained using full random genotyping and PG =2.5 and 5 was 18.3 g, nearly the same as that obtained using pedigree-BLUP (18.2 g), indicating that the former approach was not beneficial. With full random genotyping at PG = 20, the genetic gain increased to 19.7 g, about 8% more than obtained with pedigree-BLUP. Thus, when random genotyping is the only strategy used, the genotyping of a large number of candidates is vital to generate a significant genetic gain. On the other hand, genotyping combined with the use of phenotypic information produces a significantly greater genetic gain relative to that obtained with pedigree-BLUP. For example, with PG = 2.5, the dH was 19.9 g, which is 9% greater than obtained with pedigree-BLUP. With the genotyping of the heaviest 20% of candidates, the dH increased to 23.3 g, 25% greater than that obtained with pedigree-BLUP. Thus, inclusion of genomic information can substantially increase the dH with the application of a proper genotyping design.

**Table 4.**
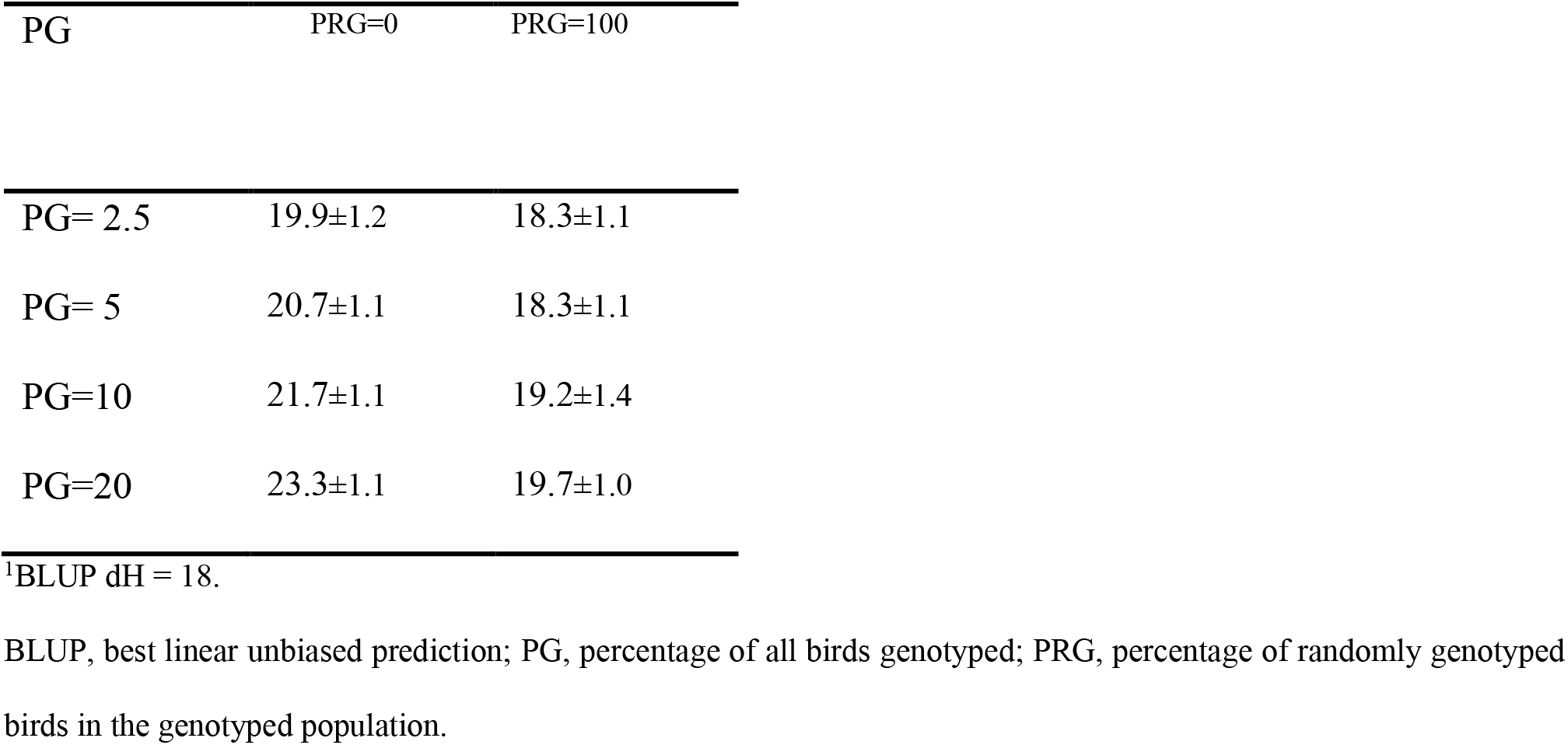
Mean (standard deviation) rates of genetic gain per time step (dH) for pedigree-BLUP^1^ *vs* selected scenarios (PG = 2.5–20, genotyping of equal percentage of both sexes randomly or with phenotypic information).

Although the incorporation of genomic information in broiler breeding programs increases the genetic gain compared with the use of pedigree-BLUP, this gain is lesser than observed in most other contexts, such as dairy cattle breeding. The genomic selection of dairy cattle was predicted to increase the genetic gain by 50–100% (Schaeffer, 2006; Pryce and Daetwyler, 2012). For laying hens, genomic prediction doubled the rate of genetic gain for egg production traits (Wolc et al., 2015) and increased the rate of genetic gain in the survival time (Alemu et al., 2016) compared with traditional pedigree-based selection. In those cases, the main benefit originated from the reduction of the generation interval. Although no such reduction occurred in this study, genomic information substantially increased the genetic gain compared with pedigree-BLUP due to increased selection accuracy. However, the genotyping strategy needs to be designed appropriately for such an effect to occur.

## Conclusion

This study was performed to investigate the importance of factors influencing the rates of genetic gain and inbreeding, to inform the design of optimal genotyping strategies for broiler breeding programs based on genomic selection and extensive phenotyping. We investigated the effects of and interactions between PG, PMG, and PRG (instead of preselection based on birds’ early phenotypic performance) using stochastic simulation with a full factorial design. All factors had significant effects, and the interaction among the three factors was highly significant. In general, the rate of genetic gain increased with increasing PG and selection of birds to be genotyped based on their early phenotypic performance (i.e., decreasing PRG). Although each factor is important, the interaction among the three factors determines the optimal genotyping strategy for a given breeding scheme. Decision making for each factor depends on the circumstances. If available resources allow for the genotyping of a small percentage (e.g., 2.5% or 5%) of all birds, genotyping of the heaviest male candidates is best. If resources allow for the genotyping of ≥40% of candidates, the genotyping of 50% of males with a small PRG (i.e., preselection based mainly on performance) is best. The incorporation of genomic information in a broiler breeding program can substantially increase the rate of genetic gain, given that a proper genotyping strategy is chosen.

## Funding and employment

SWA, ACS LW, PM, SA, and JJ were employed by Aarhus University and funded by Cobb-Vantress Inc. JH and RH were employed and funded by Cobb-Vantress Inc.

## Acknowledgments

The authors thank the genomics and quantitative genetics team at Cobb-Vantress Inc. for providing funding, as well as discussion and comments about commercial broiler breeding programs.

## Authors’ contributions

SWA designed the study and simulation, performed the simulation, and drafted the manuscript. ACS and JJ participated in the design of the study and simulation, and in the drafting of the manuscript. JH and RH participated in the design of the study and prepared the data. LW and PM helped design the study. All authors contributed to discussion of the results, and read and approved the final manuscript.

